# From Acute to Persistent Infection: Revealing Phylogenomic Variations in *Salmonella* Agona

**DOI:** 10.1101/2024.07.03.601855

**Authors:** Emma V. Waters, Winnie W. Y. Lee, Amina Ismail Ahmed, Marie-Anne Chattaway, Gemma C. Langridge

## Abstract

**Background:** *Salmonella enterica* serovar Agona (*S.* Agona) has been increasingly recognised as a prominent cause of gastroenteritis. This serovar is a strong biofilm former that can undergo genome rearrangement and enter a viable but non-culturable state whilst remaining metabolically active. Similar strategies are employed by *S.* Typhi, the cause of typhoid fever, during human infection, which are believed to assist with the transition from acute infection to chronic carriage. Here we report *S.* Agona’s ability to persist in people and examine factors that might be contributing to chronic carriage.

**Methods:** A review of 2,233 *S.* Agona isolates from UK infections (2004-2020) and associated carriage was undertaken, in which 1,155 had short-read sequencing data available. A subset of 207 isolates was selected from different stages of acute and persistent infections within individual patients. The subset underwent long-read sequencing and genome structure (GS) analysis, as well as phenotyping assays including carbon source utilisation and biofilm formation. Associations between genotypes and phenotypes were investigated to compare acute infections to those which progress to chronic.

**Results:** GS analysis revealed the conserved arrangement GS1.0 in 195 isolates, and 8 additional GSs in 12 isolates. These rearranged isolates were typically associated with early, convalescent carriage (3 weeks – 3 months). We also identified an increase in SNP variation during this period of infection.

**Conclusion:** We believe this increase in genome-scale and SNP variation reflects a population expansion after acute *S.* Agona infection, potentially reflecting an immune evasion mechanism which enables persistent infection to become established.

## INTRODUCTION

*Salmonella*, a ubiquitous Gram-negative organism, is responsible for salmonellosis, manifesting in humans as either enteric fever or gastroenteritis, caused by typhoidal and non-typhoidal *Salmonella* (NTS), respectively (1). NTS cause an estimated 93.8 million global cases per year, of which 86% are attributed to foodborne gastroenteritis (2). While NTS infections are typically self-limiting, the potential necessity for antimicrobial treatment has formidable healthcare financial implications (3).

*Salmonella enterica* serovar Agona (*S.* Agona) has been increasingly recognised as a prominent cause of foodborne gastroenteritis. In the European Union (EU), *S.* Agona has consistently featured in the top 20 serovars causing salmonellosis since reporting began in 2008 (4–9). The UK, in particular, grapples with an annual average of ∼8,500 NTS cases, of which *S.* Agona has always been among the top 10 most reported serovars (10–14). A surge in cases in 2017 raised *S.* Agona from a historical ∼1.7% prevalence (10,11) to 2.4%, sustaining this heightened level until 2019 and moving *S.* Agona into the top 5 most reported serovars in the UK (12–14). Similar trends were seen across the EU, with *S.* Agona reaching its highest-ever ranking as the sixth most common serovar in 2017 (8).

This spike in *S.* Agona cases coincided with two multi-country outbreaks under investigation in 2017 (7). One, encompassing 147 cases across five EU countries, was tentatively linked to ready-to-eat food products containing cucumbers; however, attempts to pinpoint precisely where contamination occurred in the production chain remained inconclusive (15). The other, involving 39 cases across three EU countries, was traced back to infant formula produced in a French facility, which had previously been associated with a 141-case outbreak in 2005 (16–18). Due to France’s lack of genomic surveillance pre-2017, it is unknown if these two outbreaks were caused by *S.* Agona that had persisted in the same facility for over a decade undetected. The ability of *S*. Agona to persist in dry food production environments has been documented previously. A US cereal facility suffered outbreaks separated by a decade was due to the persistence of an identical strain which was hypothesised to persist in wall cavities prior to reintroduced into the facility by a construction project (19,20), whilst Norwegian fish feed factories have seen evidence of persistence lasting at least three years due to ineffective decontamination procedures (21,22).

The ability *of Salmonella* to persist and remain viable for extended periods of time in low moisture environments is likely to be caused by many factors, which may vary between serovars and strains (23). It is well documented that switching to low moisture environments is a trigger for increased levels of osmoprotectants, rRNA degradation, filament formation, outer membrane porin transcription and alternative sigma factors (23). *Salmonella* are known for their capacity to form biofilms (24) and rearrange their genome (25), which both have the potential to enhance pathogenicity and alter phenotypic and metabolic capabilities. These survival mechanisms may also contribute to inducing a viable but non-culturable (VBNC) state, allowing persistence in food processing environments before more favourable conditions facilitate resuscitation (26).

*S.* Agona has been identified as a strong biofilm forming serovar (27) that can undergo genome rearrangement (28) and enter a VBNC state with ∼1% of the population remaining metabolically active (26). These strategies of persistence are employed by *S.* Typhi, the causative agent of typhoid fever, during human infection where it is hypothesised that genome rearrangement and biofilm establishment are required for the transition from acute infection to chronic carriage (29). The frequency and mechanism by which acute *S.* Agona infections develop into chronic carriage in humans remain largely unknown. In this study we have reviewed the surveillance data for *S.* Agona, to assess the demographic changes in the number of cases and the use of HierCC as a potential method for sentinel surveillance with respect to geographical origin per isolate. We have also investigated the genomic diversity of *S.* Agona strains that persist in the human host and assessed relevant phenotypes that might contribute towards clinical persistence.

## METHODS

### Bacterial isolates included in this study

A collection of 2233 *S*. Agona isolates referred to UKHSA’s Gastrointestinal Bacteria Reference Unit (GBRU), from local diagnostic laboratories in England and Wales, between 1 Jan 2004 and 31 Dec 2020 were included in this study. Ethical approval for the detection, identification, typing and antimicrobial resistance characterisation of gastrointestinal bacterial pathogens from infected patients submitted to the GBRU is not required as this is covered by UKHSA’s surveillance mandate. Patient consent was not required, as UKHSA has authority to handle patient data for public health monitoring and infection control under section 251 of the UK NHS Act of 2006.

For patients presenting with persistent infections (i.e. with multiple isolates), the isolation date was assessed, and carriage status was allocated as one of four categories: same episode (≤31weeks from the first submitted isolate), convalescent carriage (>31weeks –≤31months), temporary carriage (>31months – 121months) and chronic carriage (>121months) (30,31).

### Whole genome sequencing

Whole genome sequencing (WGS) was conducted as part of routine sequencing of all *Salmonella* isolates referred to UKHSA’s GBRU from 2014 onwards, as previously described (32). Prior to 2014, *Salmonella* isolates were speciated and sub-speciated using PCR (33–35). Selected strains isolated before 2014 were cultured from archived stocks prior to sequencing. Briefly, DNA was extracted using QIAsymphony DSP DNA midi kits (Qiagen) before being fragmented and tagged for multiplexing with NexteraXT DNA Preparation Kits (Illumina) and sequenced on an Illumina HiSeq 2500, yielding 100 bp paired end reads. Trimmomatic (36) was used to trim sequence data with leading and trailing at <Q30. All subsequent analysis was carried out on trimmed sequences to identify *Salmonella* species, subspecies, serovar and antimicrobial resistant determinants as previously described (34,35) FASTQ data is available from the NCBI Short Read Archive, BioProject accession PRJNA248792. Individual accession numbers for isolates analysed in this study are given in Table S1.

### Population structure of S. Agona

Raw WGS data files of 1125 isolates from cases in England were uploaded to EnteroBase (https://enterobase.warwick.ac.uk/) and were assembled using the then current backend pipelines (v3.61–4.1) including cgMLST analysis to produce a cgST using cgMLST (v2) and HierCC (v1) algorithms as previously described (37,38). There were 1078 isolates that met the cgMLST quality parameters for *Salmonella* (size >40001- ≤58001kbp, N50 ≤201kbp, number contigs ≤600, low-quality sites ≤5%, minimum taxonomic purity ≥70%) (39) and were included for analysis (Table S1). The minimum spanning tree was created in EnteroBase using the MSTree v2 algorithm and visualizing on GrapeTree (38). Previous studies have shown that analysing strains at the 5 SNP threshold might be appropriate to detect clusters or closely related clones, and that cgMLST is equivalent to SNP when detecting clusters (31,35,40–42). Therefore, HierCC was analysed at the five allelic level (HC5 – strains linked within five cgMLST alleles) for trend analysis in association with travel, year of isolation and carriage status patterns. Phylogenetic analysis was undertaken using the Ninja Neighbour Joining method (43) and visualized on iTOL (v5) (44). The historical isolates (2004-2013) were not included in the interpretation of results.

### Nanopore sequencing and hybrid assembly

A subset of 207 isolates (Table S2), comprising all viable persistent infection isolates (133) and a representative set from solely acute infection (74), were long-read sequenced at the Quadram Institute to generate hybrid assemblies with WGS data. High-molecular weight genomic DNA was extracted using Fire Monkey kits (RevoluGen) following manufacturer’s instructions. Long-read sequencing libraries, containing 48 DNA samples, were prepared using the Ligation Sequencing Kit (SQK-LSK109, Oxford Nanopore Technologies (ONT)) in conjunction with the Native Barcoding Expansion 96 Kit (EXP-NBD196, ONT). Full details of library prep are provided in supplementary information. These libraries were loaded and sequenced on MinION R9.4 flow cells (FLO-MIN106, ONT) with run times of up to 120 hours.

Raw sequencing data was collected using ONT MinKNOW software (v4.0.5) and subjected to local base-calling, de-multiplexing and barcode trimming using ONT Guppy (v5.0.11).

The open platform Galaxy (v19.05) was used to create hybrid assemblies and perform subsequent bioinformatic analysis: long-read sequences were filtered for high-quality using Filtlong (v0.2.0) (https://github.com/rrwick/Filtlong) assembled using Flye (v2.5) (45), and polished with two rounds of Racon (v1.3.1.1) (46) and one round of Medaka (v0.11.5, ONT). Hybrid assemblies were generated by using WGS data to polish the long-read assemblies with two rounds of Pilon (v1.20.1) (47). Minimap2 (v2.17) (48) was used to provide the alignment of reads to the Flye and long-read assemblies required by Racon and Pilon, respectively.

### Genomic analysis

AMR and plasmid profiles of hybrid assemblies were determined using staramr (v.0.10.0) (49) with the ResFinder (50), PlasmidFinder (51) and *Salmonella* PointFinder (52) databases, downloaded on 24 May 2022, 18 Jan 2023 and 1 Feb 2021, respectively. AMR profiles generated from these databases were checked for consistency with the UKHSA pipeline output. All discrepancies were investigated using short-read assemblies generated with Shovill (v1.1.0) (https://github.com/tseemann/shovill) and were found to be due to low coverage regions in hybrid assemblies. Virulence factor profiles of all hybrid assemblies were determined using ABRicate (v.0.9.7) (https://github.com/tseemann/abricate) with the virulence factor database (53) downloaded on 5 Nov 2021. If any differences in AMR profiles were observed between isolates from the same patient, or if a hybrid assembly had multiple genomic contigs, profiles were confirmed using short-read Shovill assemblies. Variation in biofilm and additional virulence genes responsible for salmonellosis was determined using pangenome analyses conducted with Roary (v.3.13.0) (54) from annotated files generated with Bakta (v1.6.1) (55). Additionally, amino acid substitutions within biofilm genes were identified by aligning in SnapGene software (www.snapgene.com) and their effects were predicted using the PredictProtein web server (56).

Genome structures (GSs) were identified either by *socru* (v2.2.4) (28) or by manual alignment and visualisation in Artemis Comparison Tool (v18.0.2) (57). Nucleotide variation was assessed in the 133 isolates that were associated with persistent infection using Snippy (v4.4.3) (https://github.com/tseemann/snippy) where short reads were aligned to the hybrid assembly of the earliest isolate for each patient.

### Phenotypic analysis

The biofilm abilities of the 207 isolate subset were determined using a crystal violet (CV) assay. Isolates were grown overnight in LB broth containing no additional NaCl (58) (denoted LB-NaCl). Optical density was measured with a Benchmark plus microplate spectrometer system (Bio-Rad) at 600 nm (OD_600_) to ascertain an inoculation factor.

Cultures were then 1:1000 diluted into 200 μL LB-NaCl to give an OD_600_ of ∼0.001 in 96-well polystyrene plates. After 48 hours incubation at 30 °C covered in gas-permeable seals, the wells were emptied and rinsed with water before the residual biofilms were stained for 10 mins with 200 μL 0.1% CV. The excess, unbound CV was then removed, and the wells were rinsed with water. The CV bound biofilms were then solubilised in 200 μL 70% ethanol and the absorbance was measured at 590 nm. Variations in inoculation amounts were accounted for using the inoculation factor, which were then used to correct measured CV amounts. This measurement was transformed to give a relative value in comparison to the control *S.* Typhimurium strain 14028S (accession: CP001363), which was set at 100% (59–61). The biofilm dataset was tested and found to be approximately normally distributed by the Kolmogorov-Smirnov test and through graphical visualisation. A two-sample t-test was performed to compare each carriage status group (same episode, convalescent, temporary and chronic) to the acute illness group.

Carbon source utilisation was assessed for the first and last isolates from 4 of the chronic carriers: patients 1, 4, 5 and 6. For patient 6, two additional isolates from temporary and convalescent stages of infection were also assessed. Strains were prepared from agar plates as per the manufacturer’s instructions (Biolog). PM1 panels, used with dye A, for each strain were incubated for 48 hours at 37 °C in the Omnilog (Biolog). Data were exported from the Biolog File Manager and loaded into the R package CarboLogR (62). Using the RunAnalysis() command, QC was performed and identified outlier panels that were excluded from downstream analysis. All strains had a minimum of 3 replicates that passed QC. Visualisation was performed in R (v4.2.273), using the tidyverse packages ggplot2 (v3.4.375), dplyr (v1.1.378) and gridExtra (v2.3).

## RESULTS

### Demographic overview of S. Agona infections in the UK

A total of 2233 isolates of *S.* Agona were received by UKHSA’s GBRU from England (2139) and Wales (94), between Jan 2004 and Dec 2020 inclusive. These were from 2098 patients with 105 patients having additional isolates referred after initial isolation. Patients reported to general practitioners or local hospitals with diarrhoea, abdominal pain and/or symptoms consistent with gastroenteritis. Most common specimen source as expected was faeces (2055, 92.0%), followed by urine (40, 1.8%), blood (16, 0.7%) and other sources (7, 0.3%) in which one isolate was found in the gallbladder. There were 115 isolates (5.2%) where the specimen type was not stated.

Of the 20981patients, 1035 of the patients were male (49.31%), 1054 were female (50.2%) and 9 cases were unknown, with the most common age range for infection being between 0-51years (Fig. 1A). Analysis of *S.* Agona cases referred to UKHSA’s GBRU from 2004 to 2020 demonstrates fluctuating trends in incidence rates, with notable increases observed in certain years (e.g., 2008, 2017 and 2018 with over 200 cases) and periods of decline (e.g., 2006 and 2014 with ∼80 cases, the lowest being in 2020 with 39 cases during the COVID pandemic) (Fig. 1B).

**Figure 1:**
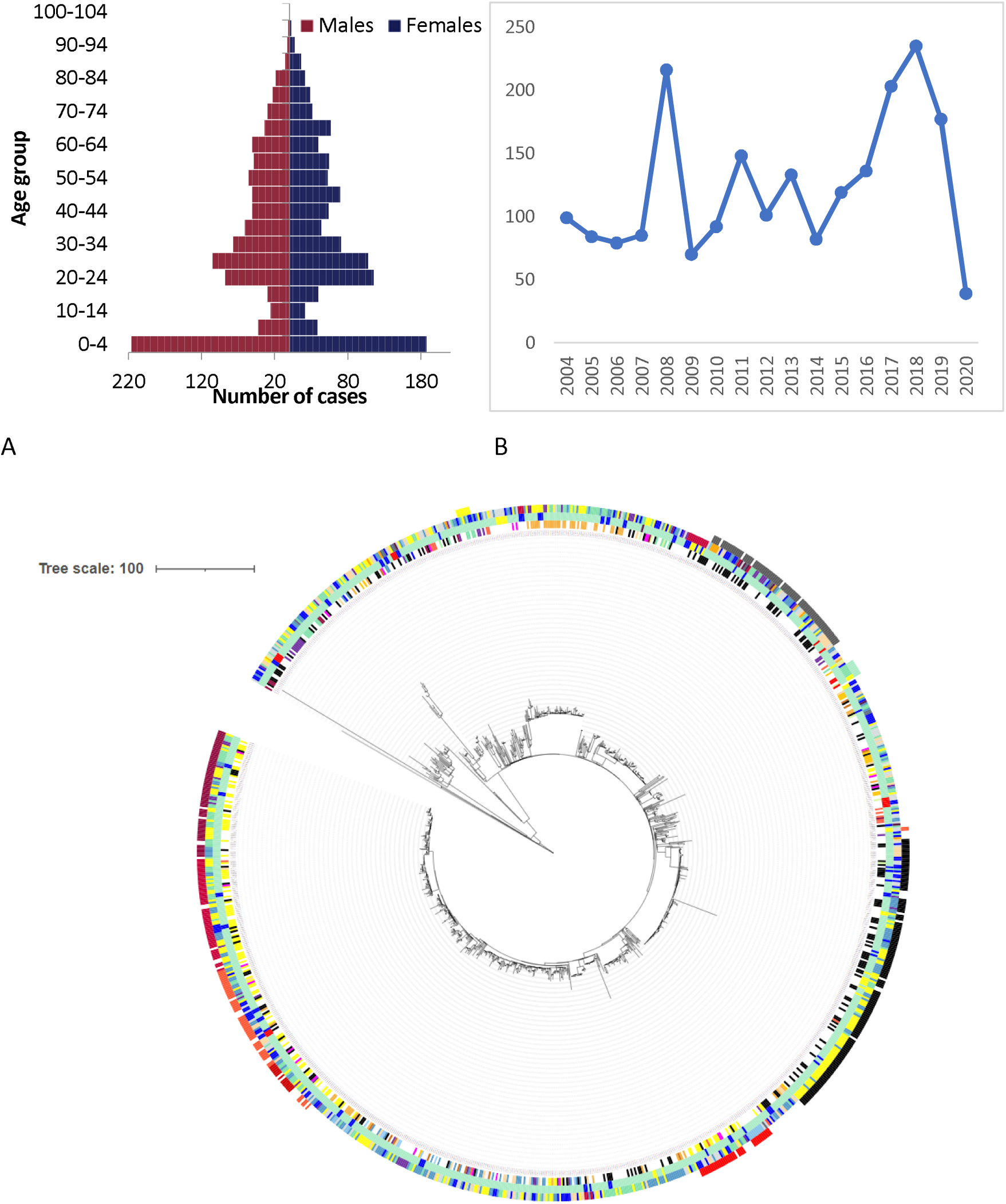
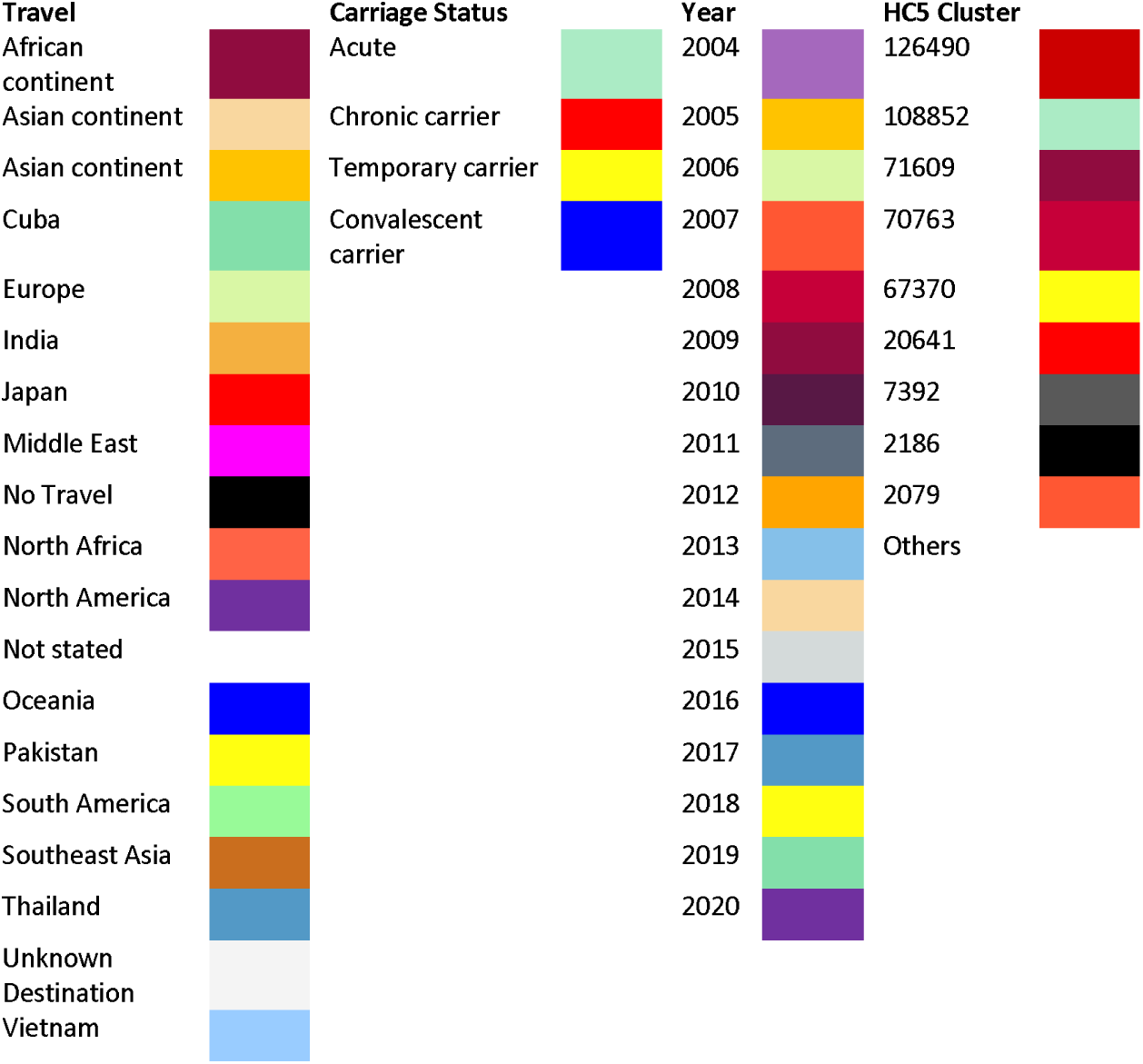
Demographic overview of S. Agona infections in England and Wales, from 2004-2020. A. Population pyramid of age and sex distribution of 2098 cases of infections. B. Annual case numbers. C. Neighbour-joining phylogeny categorised by travel (inner ring), carriage status (second ring), year of isolation (third ring) and most common HC5 clusters (outer ring) as denoted by colour. S. Agona strains associated with carriage are distributed throughout the population structure with domestic and travel associated clusters differentiated at the HC5 level.

Population structure analysis of the post genomic era (2014 onwards) show clonal expansion of strains at the HC5 level in 2017 (HC5_70668, HC5_2186) and 2018 (HC5_2186) indicating a potential outbreak may account for the increase of cases in these years though further epidemiological investigation would be needed to confirm (Fig. 1C). The majority of travel associated cases with *S.* Agona infection was from Southeast Asia with the top travel destinations being Pakistan (144, 6.9%), India (106, 5.1%), Thailand (66, 3.1%), Cuba (45, 2.1%), Egypt (38, 1.8%) and Afghanistan (34, 1.6%). Population structure analysis indicates certain clusters at the hierarchical cluster 5 level corresponds with travel to certain countries such as Pakistan (HC5_71609, 70763, 20641, 126490 & 2079), India (HC5_71231), Southeast Asia including Thailand or Vietnam (HC5_4181), Cuba (HC5_67370) and Japan (HC5_108552). There are also clonal groups not associated with travel indicating a potential domestic source in England (HC5_2186 & 7392) (Fig. 1C). Both domestic and international clusters have been around for several years showing that these strains have adapted well in their local niches and continue to thrive (Fig. 1C).

### Persistent infection exists in S. Agona

In total, 144/2233 isolates (6.4%) were attributed to persistent infection in 59 patients, which ranged from 22 days to over 6 years from first identified infection. These were categorised as follows: 86 isolates (3.9% of total collection) from 41 convalescent carriers, 36 isolates (1.6%) from 12 temporary carriers, and 22 isolates (1.0%) from 6 chronic carriers. An additional 34 isolates (1.5%) were collated from 17 patients who suffered from the same episode of acute infection (Table S1). All subsequently viable isolates associated with persistent infection (133/144) and same episode (32/34), alongside a representative set from solely acute infection (42), were long-read sequenced to generate hybrid assemblies with WGS data (Table S2).

### Strains associated with carriage are distributed throughout the population

To understand if carriage strains had any clonality, the population structure was assessed with the different carriage status of individual cases. To investigate the potential clonality of carriage strains, the population structure was evaluated based on the varying carriage statuses of individual cases. This analysis revealed that strains linked to all carriage categories were dispersed sporadically throughout the population structure. This suggests the absence of distinct clonal strains that have achieved long-term persistence and clonally expanded within the human population. Instead, it implies that numerous strains possess the capability for persistence without a single dominant clonal lineage (Fig. 1C).

### Genome analysis reveals increase in variation at transition to carriage

In accordance with the wider dataset, phylogenetic relationships between fully sequenced isolates from acute and persistent infection (Fig. S1) suggested that specific lineages were not responsible for persistent infection.

Where persistent infection did develop in a patient, we calculated the variation observed between the original isolate (first isolation) and all other isolates taken during the infection (Table S3). By adjusting for time since first isolation, we were able to determine the average number of SNPs/year identified in isolates falling within the convalescent, temporary or chronic stage of infection (Table S3). This revealed a much higher rate in the convalescent and temporary phases, averaged at 17-18 and 11-12 SNPs per year respectively. For the chronic phase the rate was 3-4 SNPs per year which is a frequency comparable to that of 1-2 SNPs per year previously reported in *S.* Agona and other NTS serovars (63,64). This raises the possibility of either hypermutation over short time span, or more likely, a larger bacterial population size being sampled from for diagnosis.

Investigating the 266 genes affected by the identified SNPs showed that 28 were observed across multiple patients. However, most of these were identified in a maximum of two patients (Table S3). Seven genes were affected in 3/4 patients: *fimD* which fimbrial outer membrane usher protein, transcriptional regulation gene *tdcA*, formate hydrogenlyase transcriptional activator *flhA*, DNA-binding transcriptional regulation gene *glpR* which responds to sugar metabolism, response regulation gene *arcA*, and *dacA* and *gyrB* which encodes for D-alanine carboxypeptidase and DNA topoisomerase respectively. Additionally, SNPs in *btuB* which encodes for TonB-dependent vitamin B12 transporter was seen in 2/6 chronic carriage patients.

We have previously shown that *S*. Agona strains typically harbour the conserved genome structure (GS1.0) but are also capable of producing extremely unbalanced GSs (28). In line with this, 94.2% (195/207) of isolates possessed the conserved structural genotype of GS1.0, which is a frequency comparable to that seen in other NTS serovars (28). However, a total of 8 additional GSs were also identified across the collection (Fig. 2A/B) and were observed only during the first 66 days of infection. Patients were considered carriers once they had harboured infecting bacteria for at least 21 days. In all bar one patient who submitted 2 samples with varied GSs, all subsequent isolates only displayed GS1.0.

**Figure 2:**
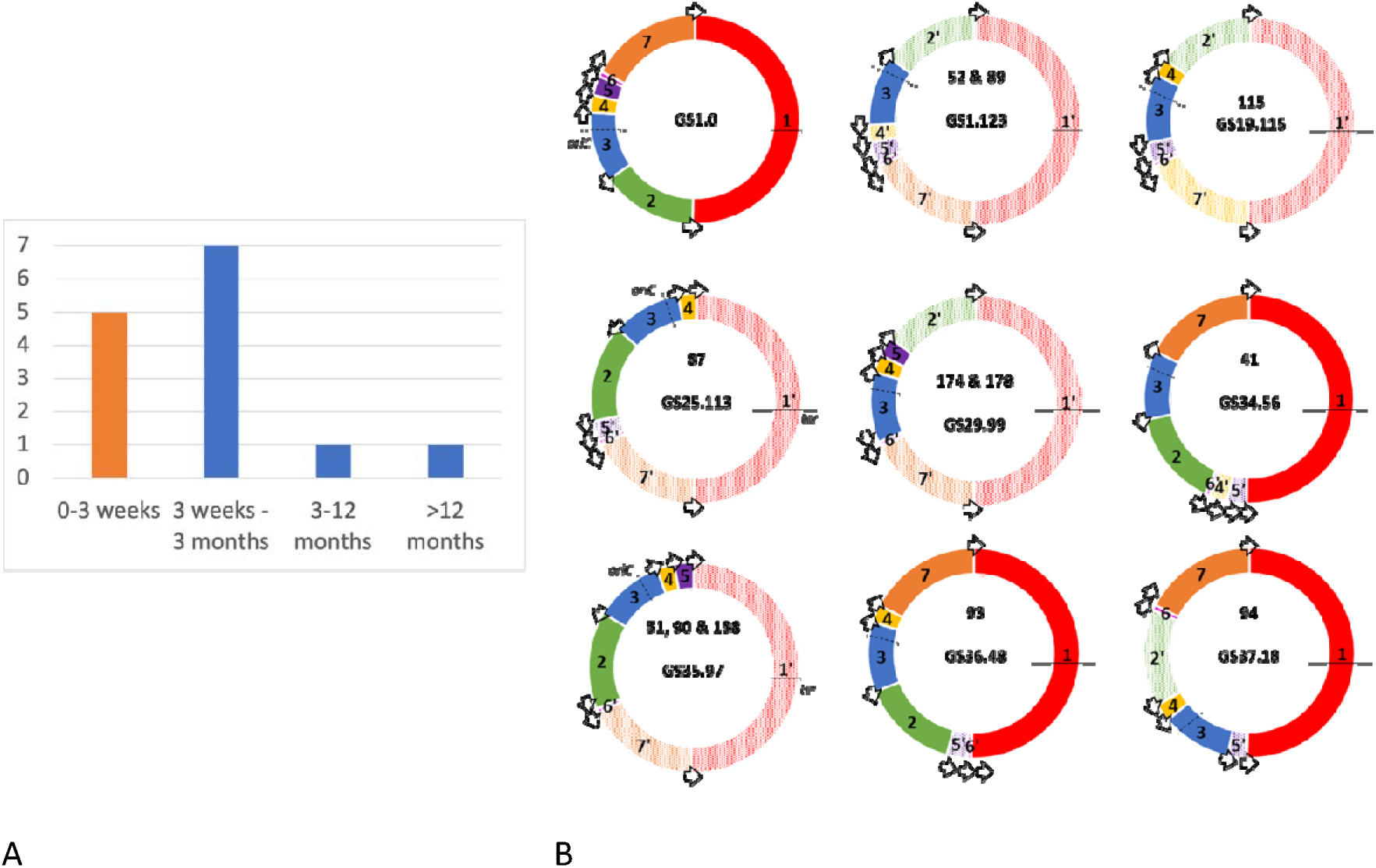
Genome structure variations were solely identified in early. A. Number and distribution of genome structures (GSs) identified from acute/same episode infection (orange) or carriage (blue). B. All GSs identified in the collection where GS designations are given as X.Y which represent the order (X) and orientation (Y) of genome fragments (coloured blocks labelled 1-7) around the chromosome (28). Isolate numbers displaying each GS are shown. Open arrows show the position of the seven ribosomal operons in Salmonella. Dashed and solid lines show the origin and terminus of replication, respectively.

### Gains and losses in metabolic capacity during infection

To identify any changes in metabolic capacity during persistent infection, carbon source utilisation was investigated by comparing the first and last isolates of chronic carriers P1, P4 and P5, spanning a timeframe of 15 to 30 months per patient (Table S4). For chronic carrier P6, carbon source utilisation was assessed longitudinally across 4 isolates spanning 16 months. Overall, metabolic capacity of isolates remained largely unchanged within each patient. However, it was striking to note that both gains and losses of metabolic ability were observed in later isolates (Fig. 3, Fig S2). The final isolates of P1 and P5 exhibited a gain in ability to utilise glyoxylic acid (P1), and L-lyxose and L-threonine (P5). Conversely in P4, the final isolate showed a gain in ability to use lactulose but a loss in the ability to use 1,2- propanediol and alpha-keto-glutaric acid. In several cases, where the initial isolate (127) of P6 was able to utilise the substrate, utilisation was much reduced in both the second (129) and third isolates (131) from the convalescent and temporary carriage phases of infection (e.g. 1,2-propanediol, alpha-keto-glutaric acid). This was even more pronounced for the third isolate (131) alone, where utilisation of many carbon sources was reduced in comparison to all other isolates, including the most recent isolate from the chronic phase of infection (Fig. 3, Fig S2).

**Figure 3:**
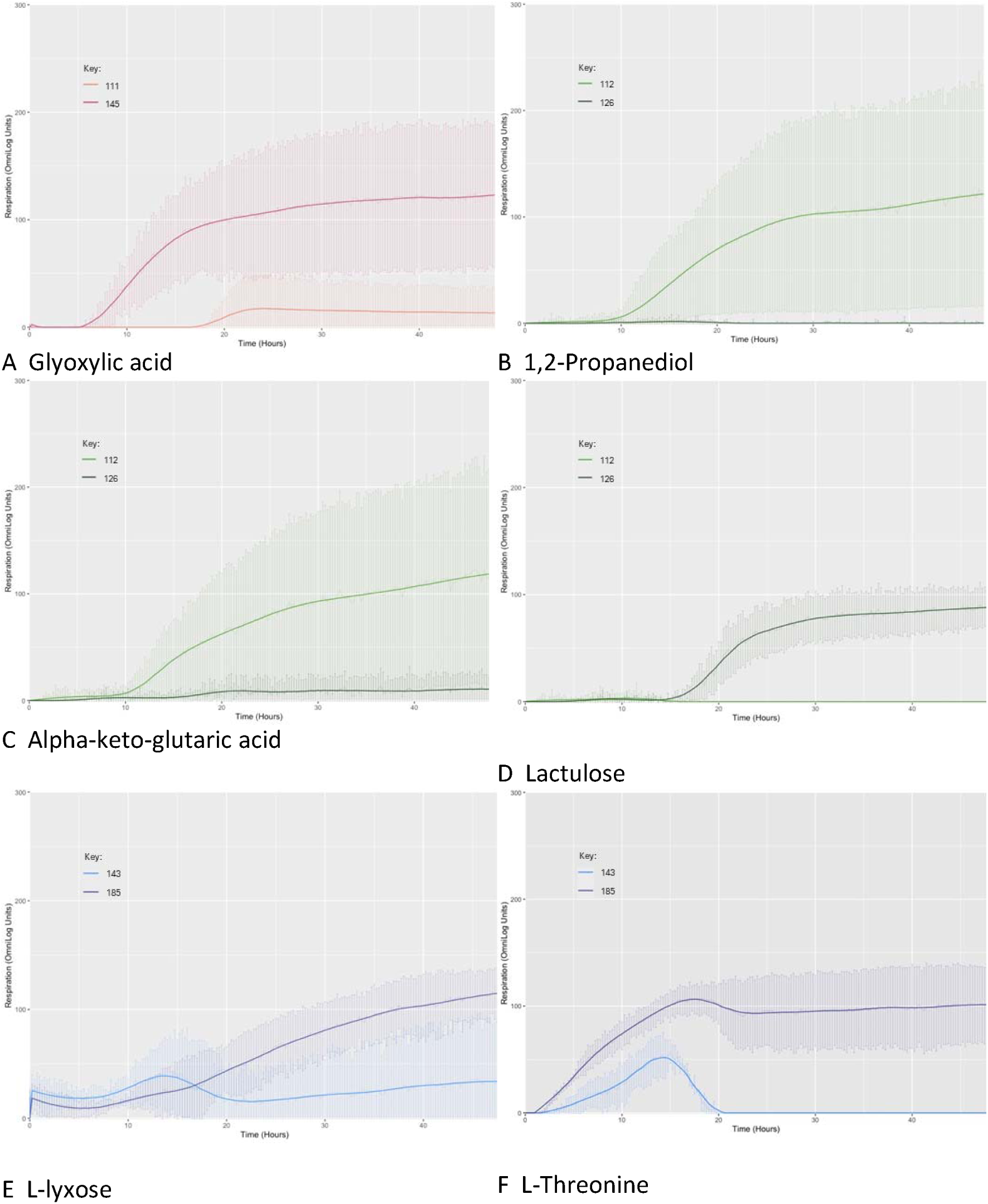
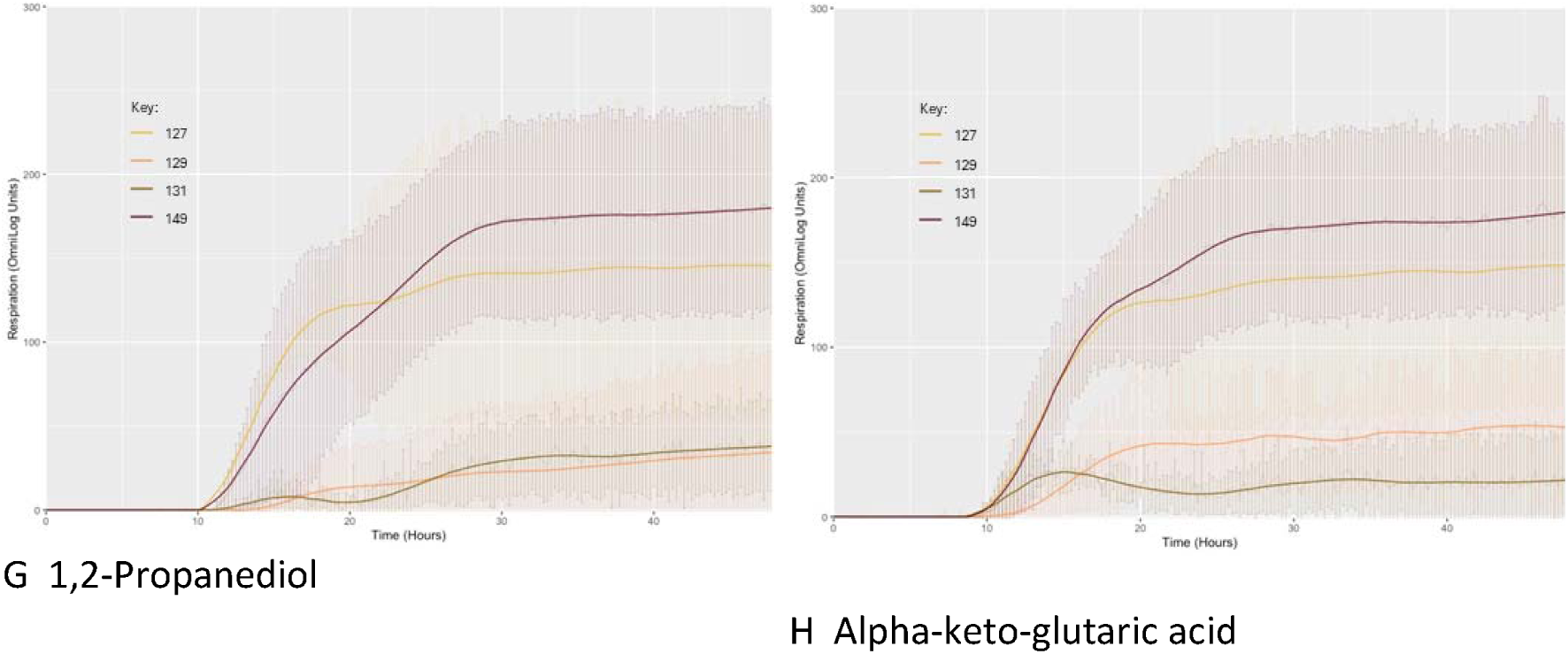
Visualisation of gain/loss in metabolic capacity during infection within patients Highlighted comparisons of carbon utilisation of original and final isolates of chronic carriers P1 (A), P4 (B-D), P5 (E and F) and P6 (G and H). For P6, carbon source utilisation was also assessed for another two intermediate isolates. Keys are included with each graph showing isolates in chronical order from the original at the top to last. Highlighted carbon source is named below each graph.

### Reduced biofilm ability associated with carriage under 12 months

The ability to form a biofilm is a key factor in S. Agona’s persistence both in and around food and has been cited in relation to disease outbreaks. We therefore assessed biofilm capacity across our collection using a crystal violet assay following growth in rich media (Fig. S3).

Biofilm formation was observed across the collection, with only 3 isolates producing no discernible biofilm at all (Table S5). We then investigated whether the biofilm ability of isolates varied in accordance with patient carriage status (Fig. S3). Our data revealed that isolates from patients with convalescent (p=0.004) and temporary carriage (p=0.002) of S. Agona had a significantly poorer ability to biofilm than isolates from patients with acute illness. To determine if this had a genetic basis, we focused on a targeted set of genes that have been previously identified as important for Salmonella biofilm formation in food production environments (Table S5) (65,66). The regulatory gene rpoS, the type III secretion system invasion gene invA and the attachment-related genes fliC and wcaA have been identified as essential/100% conserved genes in biofilm-forming Salmonella (65). These genes were identified in all isolates in this study, except in isolate 10 which lacked genes rpoS and invA.

Other genes which have been identified as having a high association with biofilm-forming Salmonella include: the additional attachment-related genes adrA, csgB, csgD, fimH and glyA; the additional regulation genes csrA, ompR and sirA; quorum sensing-related genes luxS, pfs, sdiA; and genes sipB and sipC located on Salmonella pathogenicity island 1 (SPI-1) which encode for the needle tip of the type III protein secretion system complex. The majority of these genes were detected across the entire collection. However, isolate 10 lacked both *sipB* and *sipC* genes. The lack of these genes in addition to the essential biofilm formation genes *rpoS* and *invA*, likely explains why isolate 10 did not produce a biofilm. We also looked for any amino acid changes in this gene set, identifying 10/17 as being fully unaffected (Table S5). A range of SNPs were detected that lead to synonymous substitution or changes that were predicted to have no effect on protein function (Table S5). Only 3 isolates harboured nonsynonymous changes predicted to impact function: *filC* in isolate 61, *sirA* in isolate 111 and *wcaA* in isolate 10 (Table S5).

### Plasmid presence associated with majority of resistance genes

A total of 38 antimicrobial resistance genes (ARGs), in addition to point mutations in genes encoding DNA gyrase and topoisomerase IV (*gyrA*/*parC*) were identified across 92 isolates in the collection (Table S5), associated with resistance to ampicillin, ceftriaxone, chloramphenicol, fluoroquinolones, aminoglycosides, lincomycin, rifampicin, sulfonamide, tetracycline and trimethoprim. The distribution of these resistance determinants across patients with acute and persistent infection was similar: eighteen isolates related to 38 patients with acute illness, twenty isolates to 10 patients from the same episode and fifty-four isolates from 24 patients with persistent infection. Multidrug resistant (MDR) profiles (defined as resistance to three or more antimicrobial classes) were observed in sixty-seven isolates in the collection from 37 patients, thirteen were associated with acute illness, eighteen isolates related to 9 patients with isolates from the same episode and the remaining thirty-six isolates were from 15 patients with persistent infection. AMR profiles were largely stable within patients with persistent infection (21/24 patients), with only 3 patient AMR profiles showing variable presence between isolates (Table S5).

A total of 15 different plasmid replicons were identified across 62 isolates in the collection (Table S5), including small cryptic plasmids, *Salmonella* virulence plasmids IncFII(S) and IncFIB(S) and other Inc types. Individual isolates harboured between 1 and 3 replicons each. Eleven isolates related to 11 patients with acute illness, fourteen to 7 patients with isolates from the same episode and the remainder was distributed across repeat isolates from 17 patients with persistent infection. This allowed us to investigate whether plasmid replicon presence was stable or varied within patients over time. The closest time points were from patients with isolates taken in the same episode of illness, spanning a maximum of 20 days. Within these patients, plasmid presence was largely stable (7/10 replicons), with only 3 replicons showing variable presence between isolates. In patients with persistent infection, variation was more pronounced, with 12 stable replicons within patients, and 15 showing variable presence. The variable replicons were either Col or Inc types, including the virulence plasmids IncFII(S) and IncFIB(S).

Across the 91 isolates with AMR determinants, 32 isolates were confirmed to have some or all ARGs located on plasmids. Changes in AMR profiles between isolates from persistent infection were associated with the acquisition (3/4 patients) or loss (1/4) of Inc type plasmids containing AMR determinants. Limited evidence of plasmid integration into chromosome was found in 2 acute patient isolates (#63, #64) where plasmid replicons were identified in the chromosome and in the later isolate of convalescent patient P54 (#158) where ARGs originally located on a IncX1 plasmid had integrated into the chromosome.

### All isolates have a similar capacity for virulence

A total of 154 virulence factors were identified in 207 isolates, alongside 4 additional genes that have been identified as major virulence factors responsible for salmonellosis (67–69) including: the regulation gene *hilA,* the type I secretion system gene *siiD*, the enterotoxin gene *stn*, and *fimA* which encodes for a fimbrial protein (Table S5). Demonstrating the virulence capacity of *S.* Agona, 90/158 of these genes were detected across the entire collection. Isolate 10 again lacked a high number of virulence genes in comparison to the rest of the collection, namely *hilA, sptP* and *avrA* and operons *inv, org, prg, sic, sip* and *spa. Salmonella* plasmid virulence *spv* genes and *pefA* were only found in isolates 71 and 96, corresponding to the presence of the IncFIB(S) plasmid in these isolates. No specific association between virulence factor presence and length of infection was identified.

## DISCUSSION

This research has employed routinely generated genomic data at UKHSA to gain deeper insights into the trends, prevalence and phylogenomics of *S*. Agona isolated from domestic and returning travellers in England and Wales with the view to identify and understand long term persistence in clinical infection.

While these findings can serve as a proxy for sentinel surveillance to monitor strains in different geographical regions (31) continuous real-time comparison with global data remains crucial to validate this approach and identify novel clones. In this study, HierCC at the HC5 level (39) has proven to be a valuable tool and typing scheme for assessing clonal groups across *S*. Agona strains and establishing links between population structure and case demographics (Fig. 1C). Although the study concentrates on *S*. Agona, it underscores the potential application of this approach across non-typhoidal *Salmonella* strains. Whilst *S.* Agona’s ability to persist in food processing environments and products is well documented (17,20,22), the development of *S.* Agona infections from acute into carriage is less understood. In this study we utilised a comprehensive set of 2233 *S.* Agona isolates gathered over a 16-year period from routine public health surveillance. From this collection, 6.4% of isolates were attributed to persistent infection in 59 patients, which accounted for 2.8% of the total patients in this study. This conversion percentage from acute infection to chronic carriage is similar to the 2-5 % previously seen in NTS serovars and the more well studied *S.* Typhi (30,70). The ability of *S.* Agona strains to cause persistent or acute infections does not appear to be hereditary; isolates from all carriage states were located across the phylogenetic tree demonstrating no connection between lineage and infection period.

For *S.* Typhi it is hypothesised that genome rearrangement and biofilm formation are required for the transition to chronic carriage (29); however, it was unknown if this would be the case for *S.* Agona. Our genomic analysis of a 207 isolate subset revealed that genomic rearrangement was relatively uncommon, but when it did happen, it was associated with early stages of infection. Alongside the evidence of greater SNP variation early in infection, this suggests an increase in the bacterial population during this period, plausibly required to support the transition from acute to persistent infection. An alternative explanation could be that a mixed *S.* Agona population caused these infections, which could have been present on the original food source and are now in a flux within the patient (63). It appears clear that some type of bottleneck occurs in persistent infection as variation at later stages (e.g. in temporary and chronic carriage) was lower, with only one genome arrangement observed, and very little SNP variation.

It is well established that *Salmonella* which are highly or specifically host-adapted have reduced metabolic capacity intrinsically linked to the ability to cause systemic and persistent infections. Whilst *S.* Agona is not considered to be adapted to the human host, we wanted to understand whether *S.* Agona genomes of persistent isolates displayed any hallmarks of these adaptive traits. A prime example is the inability to utilise 1,2-propanediol, a trait that is highly associated with the host restricted serovars such as *S.* Typhi, *S.* Gallinarum and *S.* Pullorum (71,72) which cause systemic, typhoid or typhoid-like disease. It has also been associated with the evolution of invasive NTS including *S*. Typhimurium ST313 (73) and *S*. Enteritidis in Africa (74). We observed this loss of function twice in our test set of 4 patients, both in isolates from carriage stages of infection and both accompanied by a SNP in *btuB*, raising the possibility that this could represent a marker of long-term persistent *S.* Agona infections.

The reduced biofilm ability associated with persistent infections of under 12 months did not appear to have a genetic basis, as almost all genes associated with *Salmonella* biofilm formation were found to be present across all isolates excluding one non-biofilm former, and the amino acid analysis indicated these were fully intact across all but 2 biofilm-forming isolates. This suggests that while these *S.* Agona strains have the capacity to form biofilms, transcriptional rewiring likely occurs during infection that reduces biofilm production, and these changes are maintained during subsequent laboratory growth.

*S.* Agona does not typically harbour IncF virulence plasmids which carry the *spv* operon and are commonly seen in *Salmonella*, but a version, *spvC*, has occasionally been found (75,76) and a megaplasmid pESI, which is predicted to provide MDR, increase virulence, and improve fitness and persistence, has started to circulate (77). Our results support that plasmid association is not a key component of *S*. Agona genomes, with only 2 acute isolates harbouring the IncFIB(S) plasmid and associated *spv* genes, but the plasmids present do attribute a considerable proportion of the AMR (42.7%) seen in this collection.

## CONCLUSION

*S.* Agona are capable of causing persistent clinical infections that can take weeks to years to resolve. Here we have identified SNP variation and changes in genome structures during early infection that may influence the transition from acute to persistent infection.

## AUTHOR CONTRIBUTIONS

GCL and MAC designed the methodology and conceived the study. EVW, WWYL and AIA performed validation, investigation and curated the data. EVW, AIA, WWYL, MAC and GCL performed formal analysis. MAC, EVW and GCL performed visualization and wrote the original draft of the manuscript. All authors reviewed and edited the manuscript.

## ACKNOWLEDGMENTS

The authors would like to thank E Holden and M Webber for providing biofilm-forming positive and negative control strains *S*. Typhimurium 14028S and 14028S ΔtolC::cat and A Nisbet for phenotype microarray visualisations. WWYL received a scholarship from the Medical Research Foundation in the National PhD Training Programme in Antimicrobial Resistance Research (MRF-145-0004-TGP-AVISO). EVW and GCL gratefully acknowledge the support of the Biotechnology and Biological Sciences Research Council (BBSRC); this research was funded by the BBSRC Institute Strategic Programmes Microbes in the Food Chain BB/R012504/1 and Microbes in Food Safety BB/X011011/1 and their constituent projects BBS/E/F/000PR10349 and BBS/E/QU/230002B.

This study is funded by the National Institute for Health Research (NIHR) Health Protection Research Unit (NIHR200892) in Genomics and Enabling Data at University of Warwick in partnership with UKHSA, in collaboration with University of Cambridge and Oxford. MAC is based at UKHSA. The views expressed are those of the author(s) and not necessarily those of the NIHR, the Department of Health and Social Care or UKHSA.

## CONFLICTS OF INTEREST

GCL has previously consulted for RevoluGen Ltd on bioinformatic analyses. Fire Monkey DNA extraction kits were provided free of charge by RevoluGen in this project.

## DATA ACCESS STATEMENT

The Illumina sequencing data generated in this study are available in DDBJ/ENA/GenBank databases under the Project accession PRJNA248792. Nanopore sequencing data and hybrid assemblies have been deposited under Project accession PRJEB76791.

## Supp Methods

For each sample, 250 ng of HMW genomic DNA was concentrated into a volume of 6.25 µL using 1X AMPure beads (A63881, Beckman Coulter), following the manufacturer’s instructions, before being end-prepped using 0.875 µL and 0.375 µL Ultra II end-prep reaction buffer and enzyme mix, respectively (E7646, New England Biosciences (NEB)) by heating at 20 °C for 5 mins and 65 °C for 5 mins. End-prepped DNA (3.75 µL) were individually barcoded using 1.25 µL of native barcode (one unique barcode for each sample), 1 µL Blunt/TA ligase master mix (M0367, NEB) and 4 µL 5X NEBNext quick ligation reaction buffer (B6058, NEB), by heating at 20 °C for 2 hours and 65 °C for 10 mins. 48 barcoded DNA samples were then pooled before being bead-cleaned using 0.6X AMPure beads with two washes of 700 µL short fragments buffer and one wash of 100 µL of 70 % ethanol before DNA was eluted in 35 µL of nuclease-free water. Sequencing adapters were ligated to 30 µL of this clean library using 5 µL Adapter Mix II alongside 5 µL T4 DNA quick ligase (E6056, NEB) and 10 µL quick ligation buffer (B6058, NEB) by incubating at room temperature for 20 mins. Adaptor-ligated DNA was bead-cleaned using a 0.6X AMPure beads with two washes of 125 µL short fragments buffer before DNA was eluted in 15 µL of elution buffer. The DNA from the last step (200–400 ng) was loaded onto an ONT MinION sequencing flow cell as directed by the manufacturer.

Supplementary Figure 1. Distribution of isolates from acute and persistent infection. Maximum likelihood tree of hybrid assemblies shows patient carriage status (key), strong biofilm formers (star) & rearranged genomes (bold text) are distributed across the tree.

Supplementary Figure 2. Comparisons of carbon utilisation of original and final isolates of chronic carriers P1, P4, P5 and P6. For P6, carbon source utilisation was also assessed for another two intermediate isolates. Keys included with each graph showing isolates in chronical order from the original at the top to last. Biolog PM1 panel carbon sources are alphabetically ordered left to right, top to bottom.

Supplementary Figure 3. Biofilm ability of isolates from acute and persistent infection. Biofilm ability is given as percentage relative to the control S. Typhimurium strain 14028S (accession: CP001363), which was set at 100% (Prouty and Gunn 2003; García et al. 2004; Trampari et al. 2019)

## Notes

### Competing Interest Statement

Gemma C Langridge has previously consulted for RevoluGen Ltd on bioinformatic analyses. Fire Monkey DNA extraction kits were provided free of charge by RevoluGen in this project.

