## Supplementary figures and images for "From Acute to Persistent Infection: Revealing Phylogenomic Variations in *Salmonella* Agona"

### Supplemental Figure S1

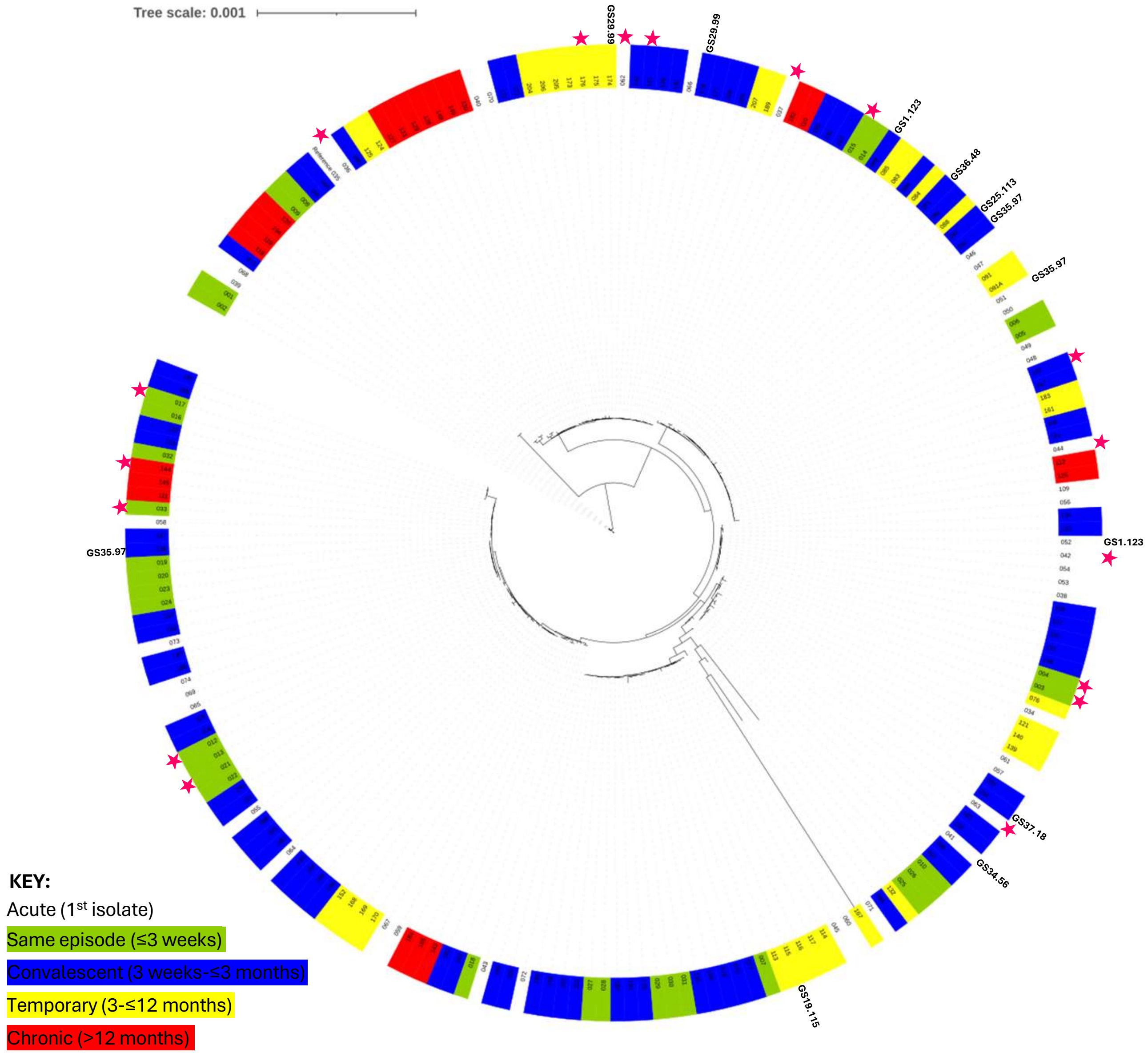

### Supplemental Figure S2

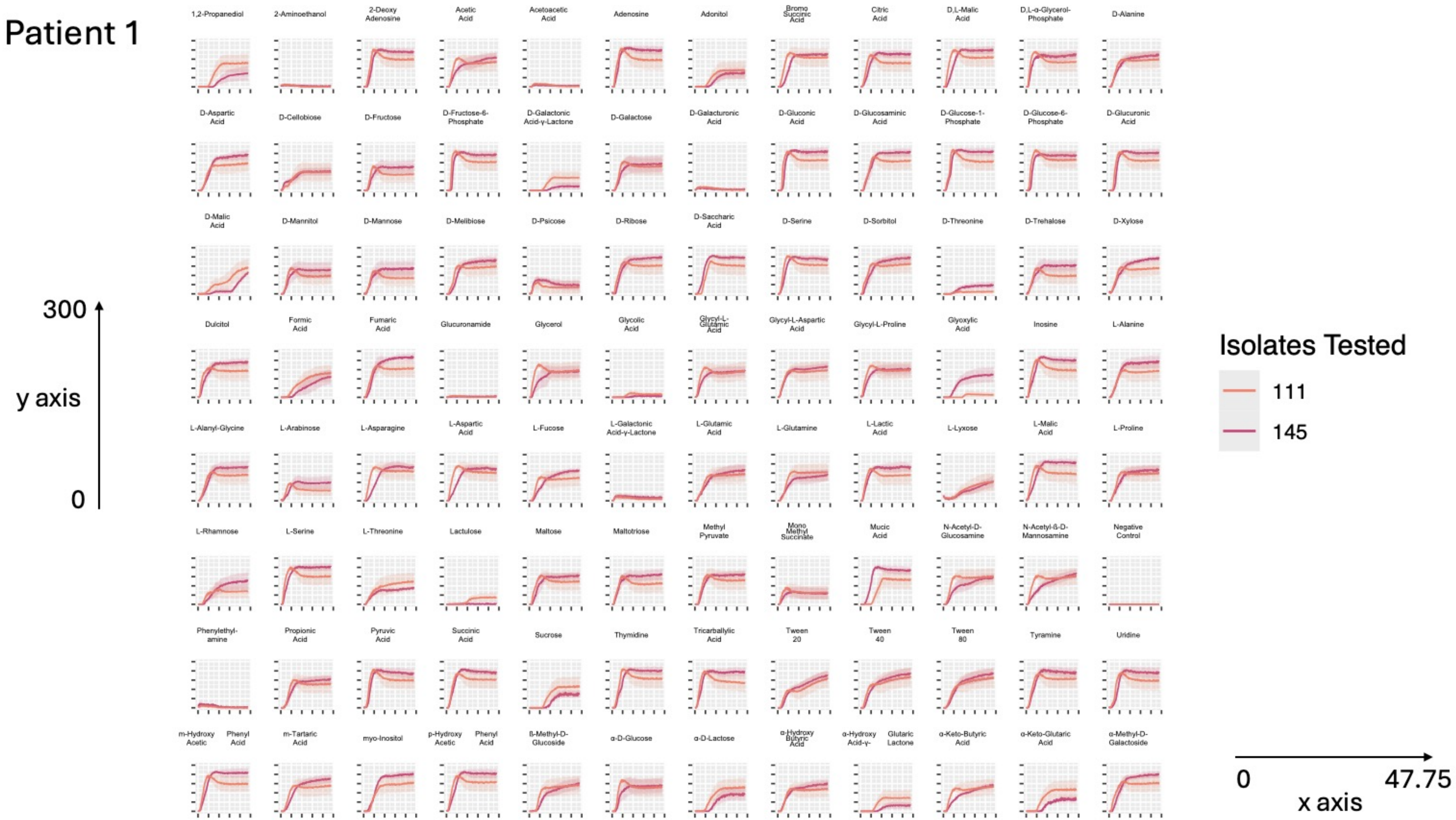

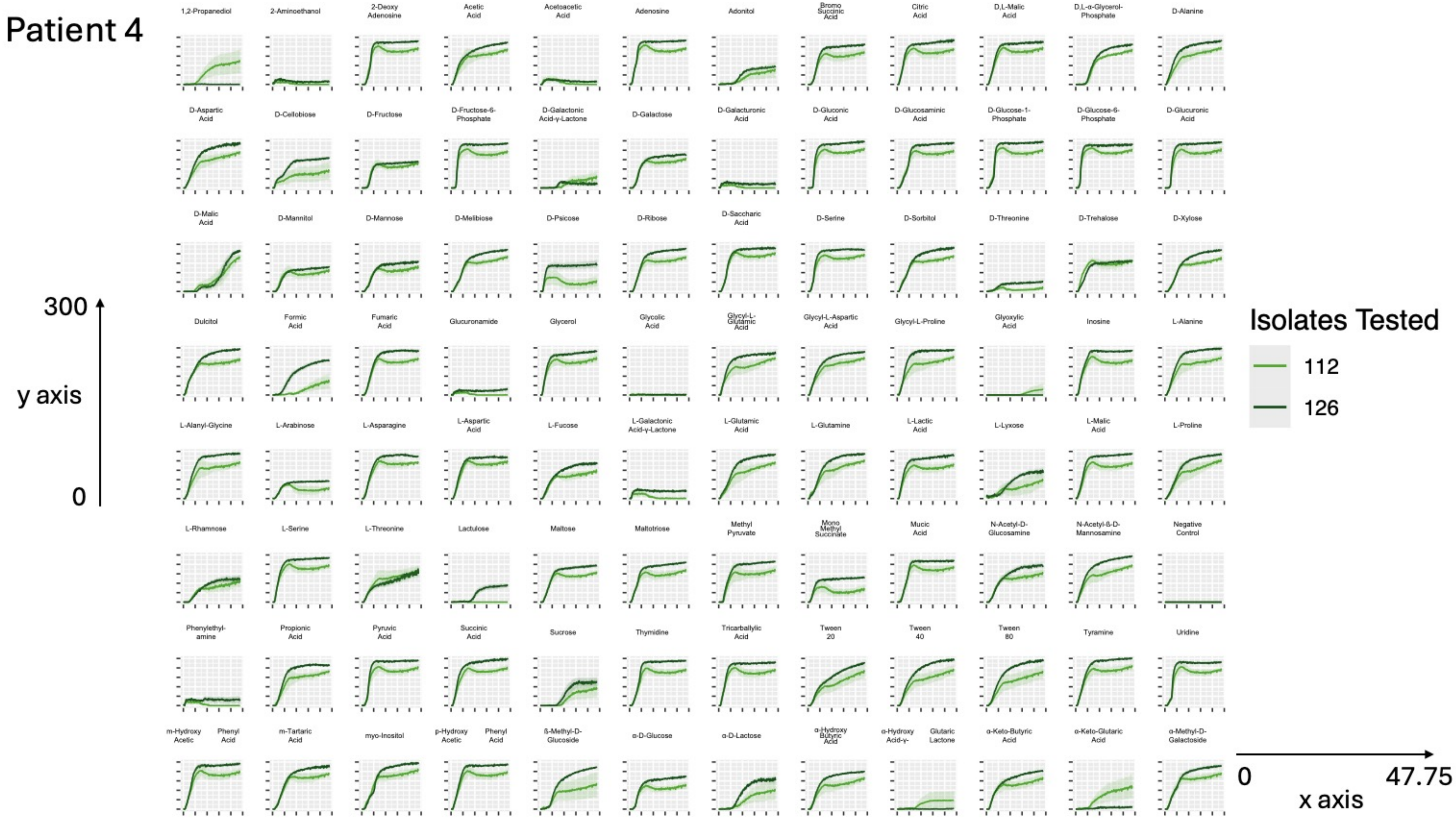

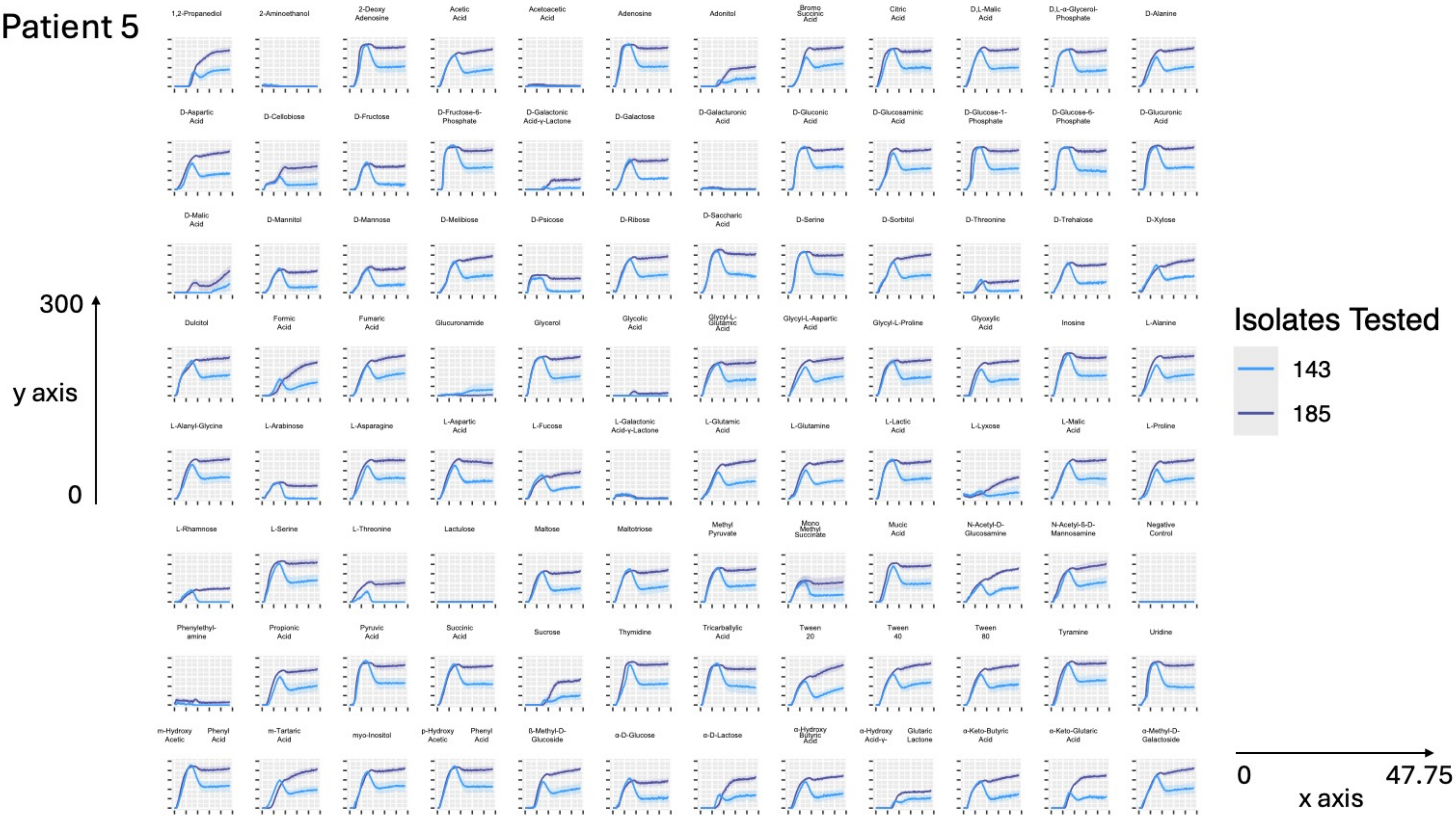

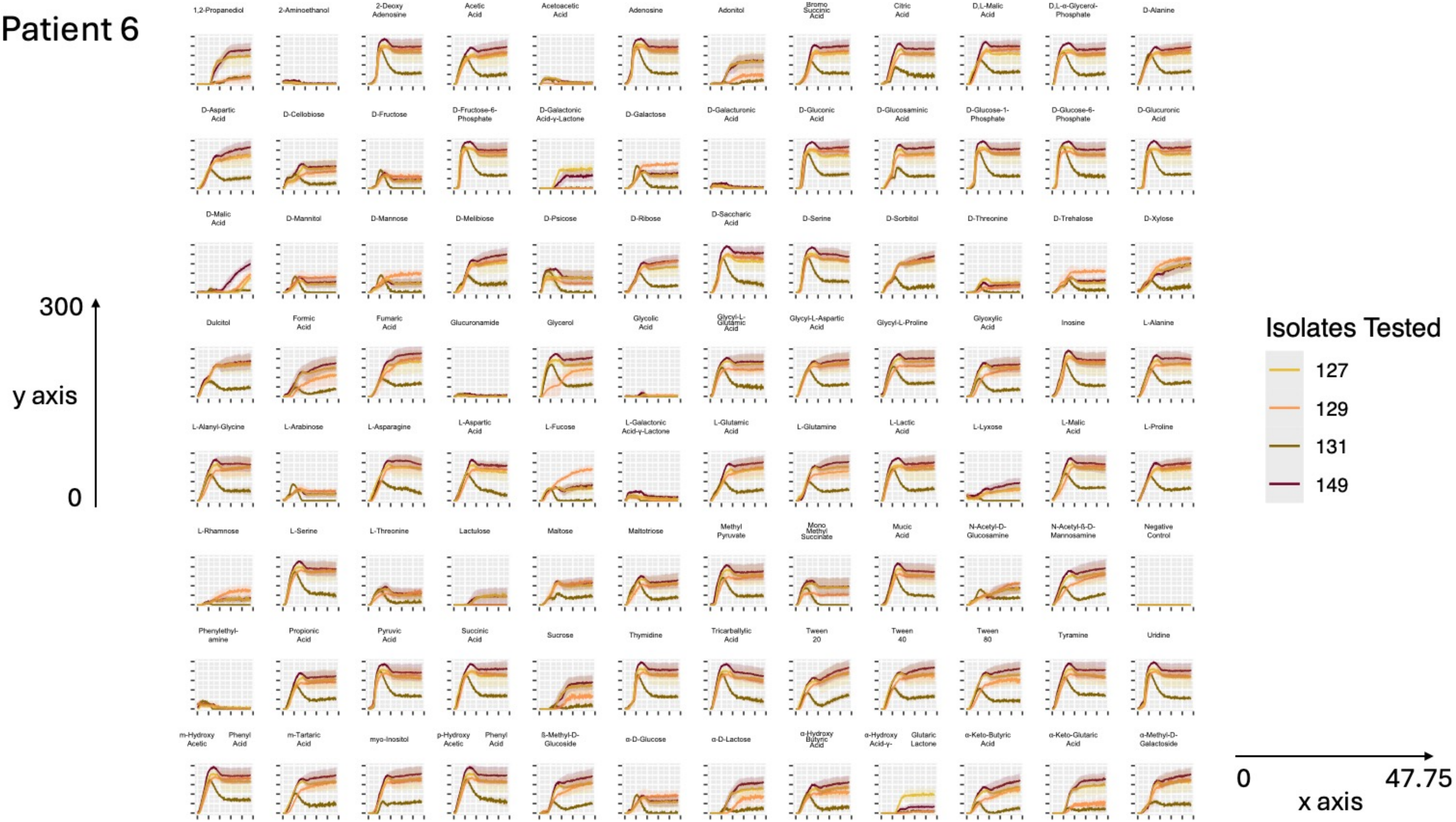

### Supplemental Figure S3

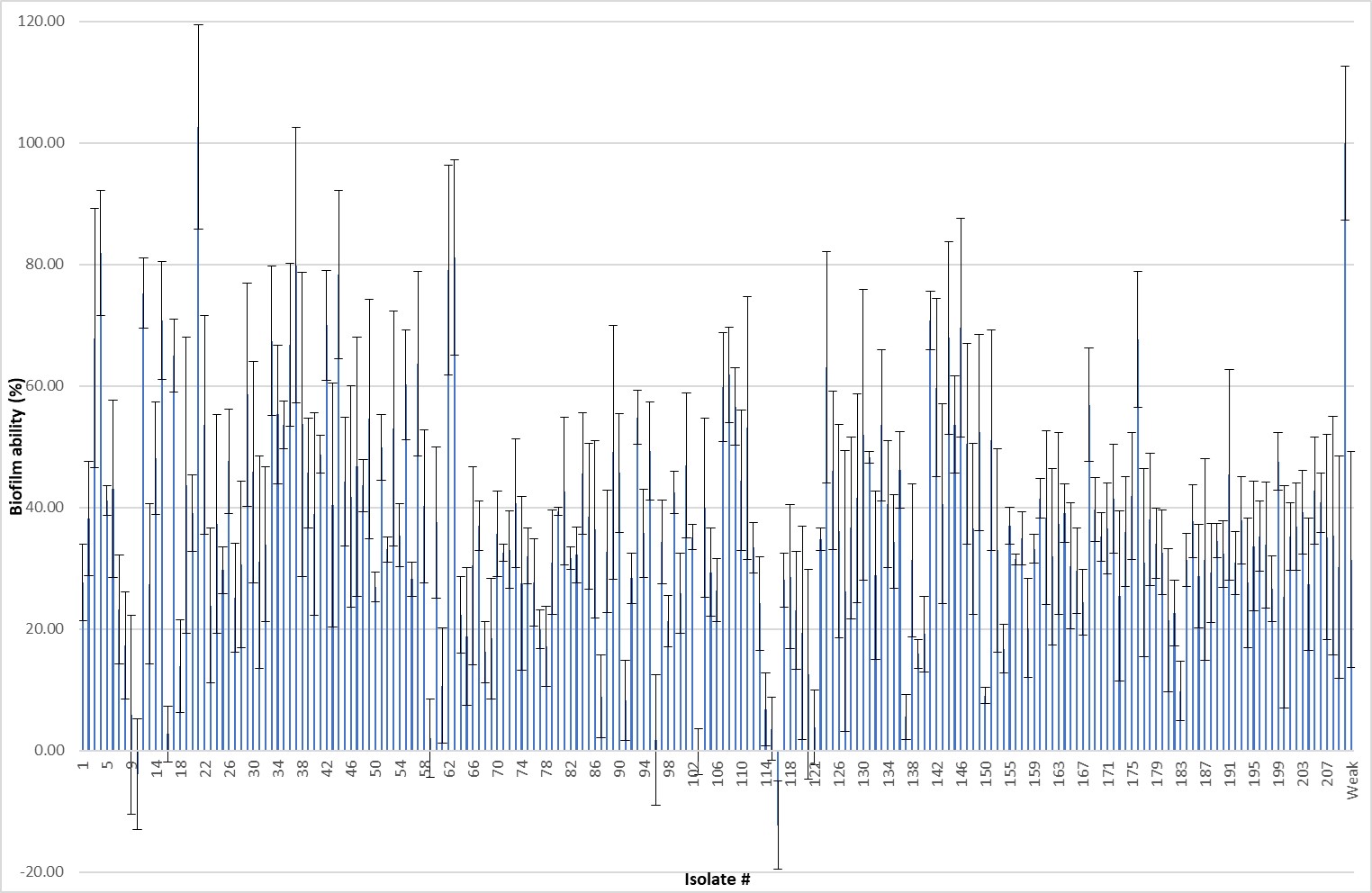
